# Pollination ecology of a threatened lupine from the core to northern edge of its geographic range

**DOI:** 10.64898/2026.04.14.718502

**Authors:** Maggie Blondeau, Cameron P. So, Anna L. Hargreaves

## Abstract

Lack of sexual reproduction limits the fitness and long-term viability of many plant populations. This may pose a particular problem for populations at the edges of species ranges, which are often small and isolated and therefore may be less likely to attract pollinators. But despite the fact that many range-edge populations are of significant conservation concern and value, there is often little information about which visitors are effective pollinators, and few explicit tests of whether range-edge populations experience reduced pollination. Here, we assess which visitors are effective pollinators of sundial lupine (*Lupinus perennis*), a legume that is threatened in much of its range, and whether pollination success varies between populations in the range core and those at the species’ northern range edge. Across six populations in the northern USA and southern Canada (Ontario), sundial lupine was visited almost exclusively by bees, but only large bees (*Bombus*, *Xylocopa*) could be confirmed as effective pollinators in single-visit experiments. While seed production varied significantly among populations, visitation rates did not. Neither pollinator visitation, pollen receipt, nor seed production declined at sundial lupine’s northern range edge. We therefore found no evidence that pollination success constrains either performance of at-risk populations of sundial lupine or the species’ northern range limit.

## Introduction

Sexual reproduction is a critical life-stage for plants, as it governs population growth rates, facilitates adaption to harsh environments, and is generally required for long-distance dispersal. For pollinator-dependent species, lack of sexual reproduction can be a particular problem in small populations, which may fail to attract sufficient pollinators (personal observations and Barrett and Kohn 1991, Ellstrand and Elam 1993, Aizen and Harder 2007). This means that pollination may be especially important for populations of species at-risk (Yakimowski and Eckert 2007) and for populations at the edges of species ranges (Hargreaves et al. 2015), both of which are predicted to be relatively small and isolated (Brown et al. 1996, Matthies et al. 2004). Indeed, sexual reproduction is the fitness component most likely to decline toward and (when measured on transplanted populations) beyond species’ range edges (Dorken and Eckert 2001, Hargreaves et al. 2014, Pironon et al. 2017).

For most flowering plants, sexual reproduction relies on animal pollinators (Ollerton et al. 2011), but whether the common declines in reproductive success across plant range limits are due to declines in pollination is unclear. Pollination success does not consistently decline toward plant range edges, though this has rarely been measured systematically (Dawson-Glass and Hargreaves 2022). Lack of pollination can play an important role in limiting ranges of some plant species (Chalcoff et al. 2012, Moeller et al. 2012, Benning and Moeller 2019) but is relatively unimportant for others (Hargreaves et al. 2015, Theobald et al. 2016, Macrì et al. 2021). This knowledge gap is particularly important for the many range-edge populations that are also of high conservation concern (Berg et al. 1994, Hardouin and Hargreaves 2023). For many at-risk plants we lack basic information about which flower visitors are effective pollinators, whether visitation rates vary among populations, and whether pollination limits reproduction (MacPhail et al. 2018, Caissy et al. 2020).

Here, we use observations and experiments to assess which floral visitors are effective pollinators of a threatened perennial plant, sundial lupine (*Lupinus perennis*, Fabaceae; Fig.1), and whether pollinator service declines from the species’ range core to its northern range edge. Sundial lupine is the only legume native to the northern eastern USA and southeastern Canada, and produces striking floral displays, meaning its occurrence and range boundaries are well-documented. Sundial lupine requires pollinators to produce seeds (Shi et al. 2005) and out-crossed flowers make twice as many seeds as self-pollinated flowers (Michaels et al. 2008). In the range core, sundial lupine is typically visited by both large and small bees, which have been observed to contact anthers and stigmas (Bernhardt et al. 2008). However, which visitors effectively transfer pollen has not been experimentally confirmed, and pollination studies have so far been limited to populations in the species’ range core. Thus it is unknown whether declining pollination limits performance at the species’ range edge, potentially reinforcing the current range limit and impeding conservation of northern, at-risk populations.

**Figure 1.**
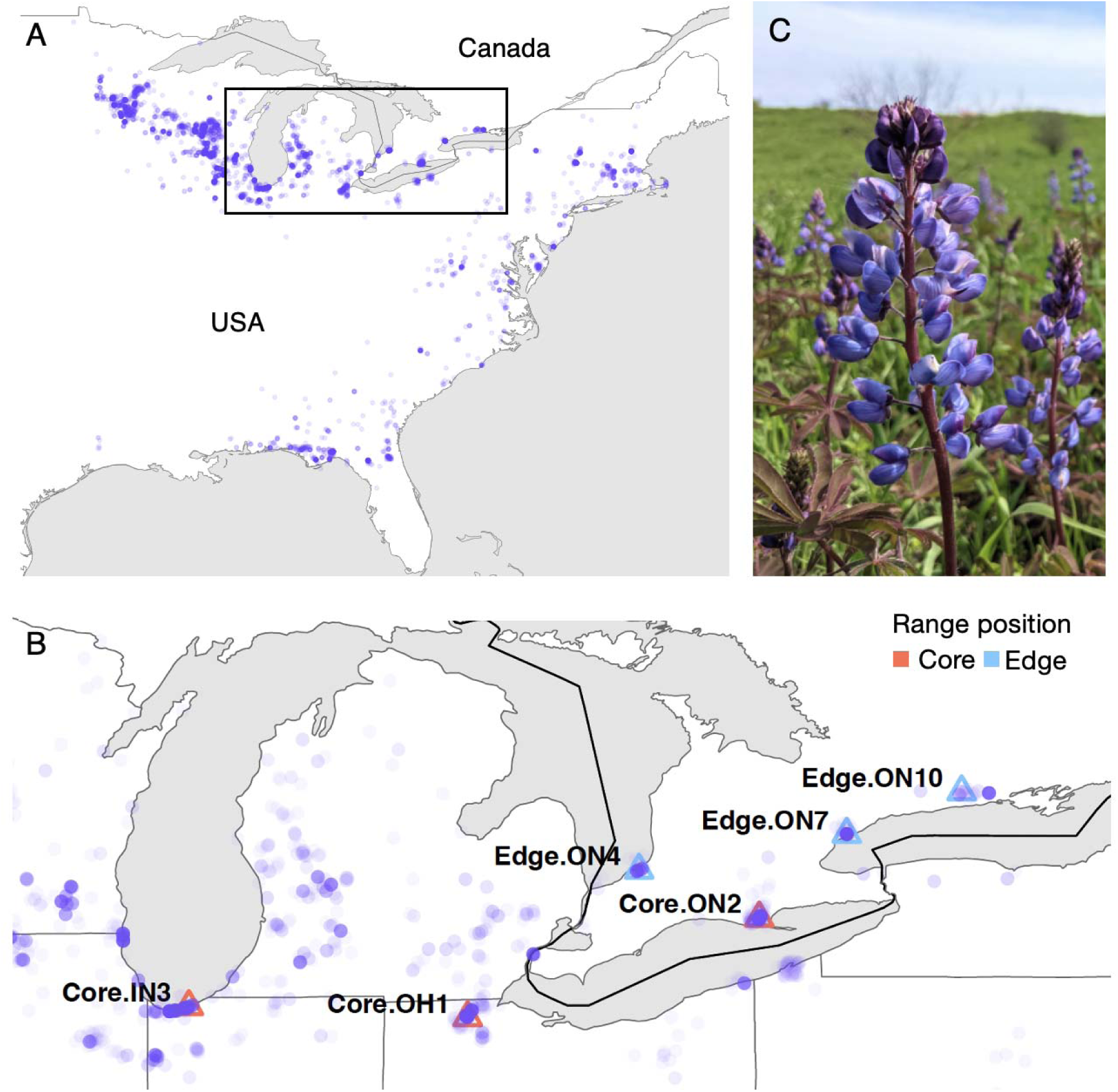
Geographic range, study sites, and flowers of sundial lupine. **A**) All research-grade observations from iNaturalist (downloaded December 6, 2023). Rectangle shows study region in B. **B**) Locations of our study populations. Population names denote each populations’ range position (range core, in red, or range edge, in blue), state or province (IN=Indiana: OH=Ohio; ON=Ontario), and a unique number within each jurisdiction x range position combination (numbers are not sequential as they are from another study with more populations). We surveyed pollinator visitation and pollen receipt in all six populations, and studied pollinator effectiveness in Core.ON2. **C**) A blooming inflorescence of sundial lupine.

We studied pollination ecology in 3 populations in sundial lupine’s range core and 3 populations at its northern range edge (Fig. 1) to answer three questions. 1) Which flower visitors are effective pollinators? 2) Do visitation rates and/or pollen receipt decline toward the species’ northern range edge? 3) Does seed production decline from core to edge populations, or co-vary with visitation rates?

## Materials and Methods

### Study species and location

Sundial lupine is associated with sandy soils of dunes and oak savannahs (Nuzzo 1986). Its range is shaped like a ‘7’, sweeping west-to-east along the Great Lakes, where its overlaps Canada in the southwestern tip of Ontario, then south along the Appalachian Mountains (Fig. 1A). Sundial lupine is threatened through much of its range, particularly at its northern range limit in Canada where it is considered nationally imperilled (NatureServe 2023, Petitta et al. 2024). We studied six populations in the Great Lakes region of sundial lupine’s range (Fig. 1; Table S1). Populations were defined as local groupings separated from other sundial lupines by at least 100 m and that experience a common management regime. The 3 ‘range-core’ populations have many populations surrounding and between them, whereas the 3 ‘range-edge’ populations are each the northernmost population at their longitude and are isolated from other lupine populations.

Sundial lupine (hereafter ‘lupine’) reproduces sexually and asexually, and sexual reproduction requires pollinators (Shi et al. 2005). In our study area, populations bloom from May to July, and individual flowers remain open for 3 to 5 days. Flowers are clustered into tall spiked inflorescences (racemes; Fig.1C), almost always 1 inflorescence per rosette (0.02% of 519 of rosettes had 2 inflorescences; Blondeau 2023)). Each flower has a banner petal, two wing petals, a keel petal that encloses the anthers and stigma (Fig. S1), and one ovary with 5 to 7 ovules (Shi et al. 2005). Flowers produce little if any nectar (we were unable to detect any). Flowers open starting at the bottom of the inflorescence. Flowers are partially protandrous; pollen is present upon opening and remains viable for 5 days, while the stigma is receptive from day 2 to 8 (Shi et al. 2005). However, strong herkogamy prevents autonomous self-pollination (Michaels 1995, Shi et al. 2005). Fruits are elongated pods, which mature from mid-June to July and disperse seeds ballistically up to 5 m (Trudeau and Framer 1996). Asexual reproduction occurs underground via rhizome spread and subsequent ramet growth.

### Pollinator observations

We surveyed floral visitation rates in 2022. In each population we did at least 10, but normally 15, observation periods over the span of three days. We surveyed visitors during high-activity periods, i.e. between 10h and 16h, low wind, full sun, and warm (18 to 28 °C). For each observation period, we selected a 1 m^2^ plot of flowering lupines in a dense area of the population and recorded the number of inflorescences and open flowers within. For 10–20 min, a single observer recorded all flower visitors (any animal interacting with the inside of a lupine flower), including the number of flowers and inflorescences each animal visited in the plot, and then identified the visitor to taxonomic Family, and to genus when possible (not always possible for small bees as we were not catching visitors in 2022). We observed how each visitor handled flowers and whether it contacted anthers and stigmas, although these are often enclosed within petals and hard to observe directly.

### Pollinator effectiveness

In 2023, we spent 2 weeks in a range-core population (Core.ON2) to assess which visitors were effective pollinators. First, we assessed which visitors carried lupine pollen. We opportunistically caught insects seen visiting lupine flowers, chilled them overnight to slow their movements, then swabbed them with fuchsin jelly (gelatin stained with basic fuchsin) to pick up externally carried pollen (Beattie 1971). For bees, we swabbed over corbiculae but never included pollen packed into corbiculae, as this pollen is generally unavailable for pollination. For butterflies, we confirmed visually that individuals were not potentially one of the at-risk species in the area before catching them, and took care not to touch butterfly wings when swabbing for pollen. We melted fuchsin swabs onto microscope slides, and later counted the stained pollen grains under 100x magnification. We distinguished lupine pollen by its characteristic shape (sundial lupine was the only lupine species at any of our populations). We euthanized one voucher specimen per bee species, storing them in 70% ethanol for identification; all other visitors were photographed and released.

Second, we confirmed which insects effectively transfer lupine pollen to stigmas using single-visit experiments (Spears Jr 1983, Hargreaves et al. 2008). We identified inflorescences with flowers whose stigmas would soon become receptive, and isolated them from visitors using 40 x 30 cm white tulle-mesh bags (Fig. S2). Once an inflorescence had receptive flowers, we removed the bag and watched the virgin, receptive flowers until an insect visited at least one of them. Insects were allowed to visit as many flowers on the inflorescence as they wanted. We then identified the insect, marked which virgin flowers had been visited (permanent marker dot at the base of the pedicle), and re-bagged the whole inflorescence to prevent further visits. We collected the stigma the next day, snipping the style 0.5 cm below the stigma and storing it in 70% ethanol. We later mounted each stigma on a microscope slide in fuchsin jelly, and counted the pollen grains adhering to the stigma. We repeated this experiment on 15 inflorescences, but due to low visitation rates only 6 of these were visited during experiments.

### Pollen receipt

In 2023, we examined pollen deposition through time and space by counting the lupine pollen deposited on stigmas collected from flowers at the end of their female phase (when stigma is elongated and ovaries are swollen). To examine temporal variation in pollen deposition, in population Core.ON2 we collected stigmas every 3 d over 2 weeks during peak flowering (30 stigmas/day; two from 15 rosettes). To examine spatial variation, we collected stigmas from the other five populations once during peak flowering. At each population, we collected two stigmas from 30 rosettes separated by ≥5 m to avoid sampling ramets of the same clone, though we reduced the distance to ≥1 m in our two smallest populations (Edge.ON 4 and 7; Table S1, total = 60 stigmas per population). Stigmas were stored in 70% ethanol, then mounted on microscope slides in fuchsin jelly. We counted the lupine pollen grains adhering to each stigma (microscope 100x).

### Seed production

In 2022 we revisited sites once plants had begun maturing seeds and surveyed reproductive success at 30 randomly-chosen rosettes per population (61 plants at Core.ON2 due to additional sampling for another study). To avoid sampling ramets of the same clone, we chose rosettes at least 5 m apart, except in our two smallest populations (Table S1), where minimum separation was 2 m. We recorded whether each rosette was reproductive; if it was we counted the fruits/inflorescence and the viable seeds in 5 randomly-selected fruits. Seeds were deemed viable if they were full and firm, and inviable if they were shrivelled and discoloured, or underdeveloped and soft (subsequent experiments found a ∼95% germination rate of seeds deemed ‘viable’). We estimated seed production per flowering rosette as: mean viable seeds per fruit x # fruits.

### Pollen limitation and breeding system experiments

We attempted to test for site differences in pollen limitation and self-compatibility by adding outcross pollen to open-pollinated flowers, and outcross- or self-pollen to virgin flowers in each population (methods in Supplementary Material). After hand pollinations we bagged each experimental flower individually to ensure we could count seeds if pods exploded (Fig. S2). However, despite being gentle while securing bags, all bagged flowers aborted.

### Statistical analyses

Analyses were conducted in R v.4.3.1 (R Core Team, 2023). Data and code will be publicly archived. We used the package ‘glmmTMB’ for model fitting (Brooks et al. 2017) and ‘DHARMA’ for model diagnostics (Hartig 2022). We assessed the significance of predictors (including interactions) by comparing models with and without the predictor of interest using likelihood ratio tests compared to a Chi-squared distribution. When a response differed significantly among sites, we tested which sites differed from each other using Tukey corrected post-hoc comparisons using the package ‘emmeans’ (Lenth et al. 2025).

#### Pollinator effectiveness

We tested whether large and small bees carried different quantities of lupine pollen using a generalized linear model (GLM) with a negative binomial error distribution (logit link) and bee type (large or small) as a predictor. Pollen deposition during single-visit experiments is described in the Results but not analysed statistically due to low sample size.

#### Visitation rates

We tested whether visitation rates in 2022 (visits per flower per minute) varied geographically. We first tested whether overall visitation rates (all bees) differed among our six sites (GLM) or between range positions using a generalized linear mixed model (GLMM) with a random effect for site. Because there were many observation periods without visits, visitation rates were zero-inflated and often <1 (i.e. non-integer data); GLMMs therefore used a tweedie error distribution (log link function). We then reran analyses including the predictor ‘bee type’ (large or small) and its interaction with site (with a random effect for observation period) to test whether geographic patterns in visitation rates varied between large and small bees.

#### Pollen receipt

We analysed pollen receipt in 2023 (lupine grains / stigma) using zero-inflated negative binomial GLMMs. We tested whether pollen receipt changed across time in population Core.ON2 using a GLMM with collection date as a categorical fixed predictor and (n = 172 stigmas total). We tested whether pollen receipt varied among sites using GLMM with site as a fixed predictor (n = 258 stigmas total). Both models included rosetteID as a random effect to account for two stigmas coming from the same rosette. We did not test whether pollen receipt was predicted by visitation rate as these data were collected in different years.

#### Seed production

We tested whether seeds per flowering rosette in 2022 varied between range positions, among sites, and with 2022 visitation rates by all bees or by large bees specifically. Models of site and range position used a negative binomial GLM and included a *ziformula ∼1* to account for zero-inflation, as many flowering rosettes failed to make fruits, and the range-position model included a random effect for site. As seed production and visitation rates were measured on different plants, these analyses used the mean visitation rate and mean seeds/rosette for each site (n = 6 data points; data were not zero-inflated).

## Results

### Effective pollinators

Lupine flowers were visited almost exclusively by bees. During 18 h of systematic observation in 2022 plus ∼15 h of opportunistic observation in 2023 (during single visit experiments and while catching visitors to assess pollen loads), we saw a total of 210 large bees (*Bombus, Xylocopa, Apis, Andrena*), 162 small bees (*Lasioglossum, Augochlorella, Hoplitis*), and 4 butterflies feed from lupine flowers.

Visitors manipulated lupine flowers differently, potentially differing in pollination effectiveness (Fig. 2). Large bees manipulated flowers’ keel petal with their mouthparts to extract pollen. They shifted the keel downwards, resulting in pollen being expelled and the stigma being extruded through the tip of the keel and contacting the bees’ thorax. The largest bees (genera *Bombus, Xylocopa*) were heavy enough that they also pushed the wings of the flower backwards the keel fully downwards to completely expose the pollen and stigma, and almost always contacted the stigma. In contrast, small bees crawled into the wing petals and only sometimes into the keel, so rarely contacted reproductive parts (Fig. 2). *Bombus* and *Xylocopa* bees tended to land on flowers at the bottom of the inflorescence (i.e. female-phase flowers) and work their way up, often visiting all open flowers before moving to a new inflorescence. Other bees did not visit flowers in any particular order, and often visited only a few flowers of the open flowers on an inflorescence. Butterflies landed on flowers and may have probed briefly for nectar (this was hard to observe with certainty) but never visited multiple flowers and are unlikely to have contacted stigmas.

**Figure 2.**
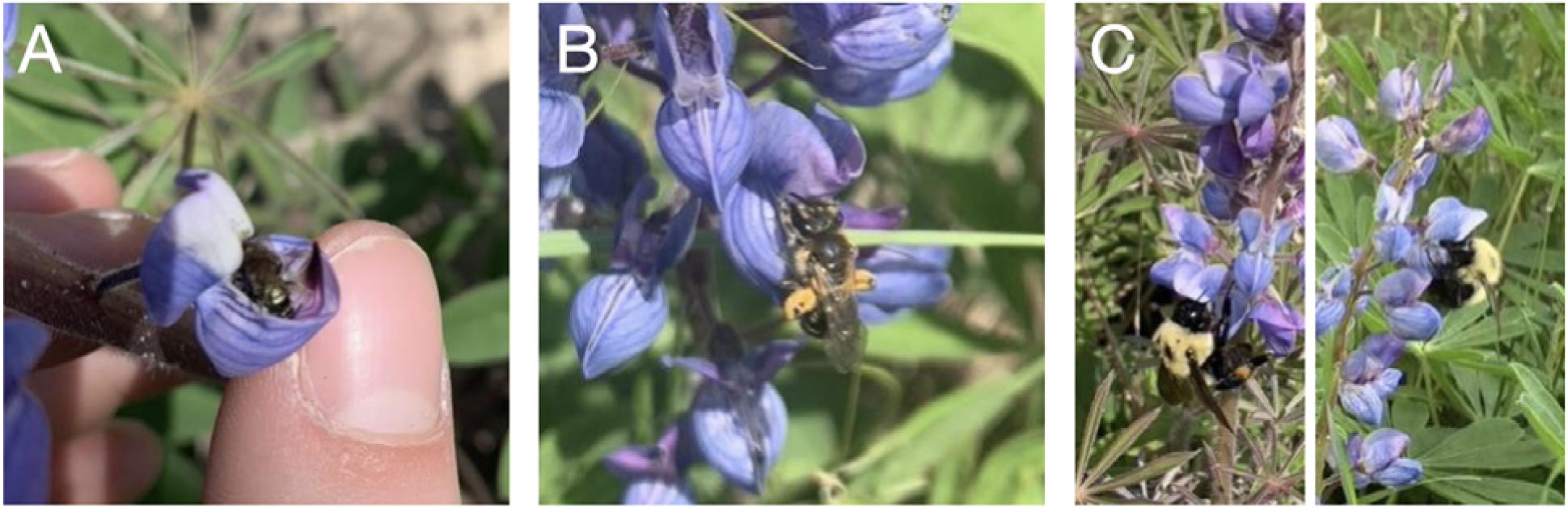
Bees visiting flowers of sundial lupine. A) ‘Small’ bee gathering pollen. The bee manipulated the keel petals while inside the wing petals, but does not interact with the tip of the keel where the stigma is housed, and so did not contact the stigma. B) ‘Large’ bee gathering pollen, manipulating the keel while weighing the flower down. While this bee was not heavy enough to push wing petals backwards, it still shifts the keel downwards to expose some pollen and the stigma, and the bee contacted the stigma. C) ‘Large’ bees (*Bombus*) gathering pollen, manipulating the keel while weighing down the flower. These bees were heavy enough to push the keel downwards to completely expose the pollen and stigma, and contacted the stigmas.

More detailed studies at population Core.ON2 confirmed that insects differed in their effectiveness as lupine pollinators. Overall, bee visitors carried from 0 to 4778 lupine pollen grains (n = 67 bees; Fig. 3A). Large bees carried more lupine pollen on average than small bees but the difference was not significant (χ^2^ = 0.18, *df* = 1, *P =* 0.68; Table S2). Only three butterflies visited lupine flowers, and carried 1, 3 and 12 lupine pollen grains respectively. We captured 15 single-visits to virgin flowers, by 4 large bees and 2 small bees. During 13 flower visits by large bees (*Bombus* and *Xylocopa*), bees deposited 1 to 67 lupine pollen grains per stigma (mean = 26.7 ± 6.2), and 12 of 13 such visits deposited at least 7 grains (i.e. at least as many as the maximum number of ovules in sundial lupine flowers; Table 1). We captured only two visits to virgin flowers by small bees (*Lassioglossum*), in which one bee deposited no pollen and the other deposited one lupine pollen grain. Thus, only large bees could be confirmed as effective pollinators.

**Figure 3.**
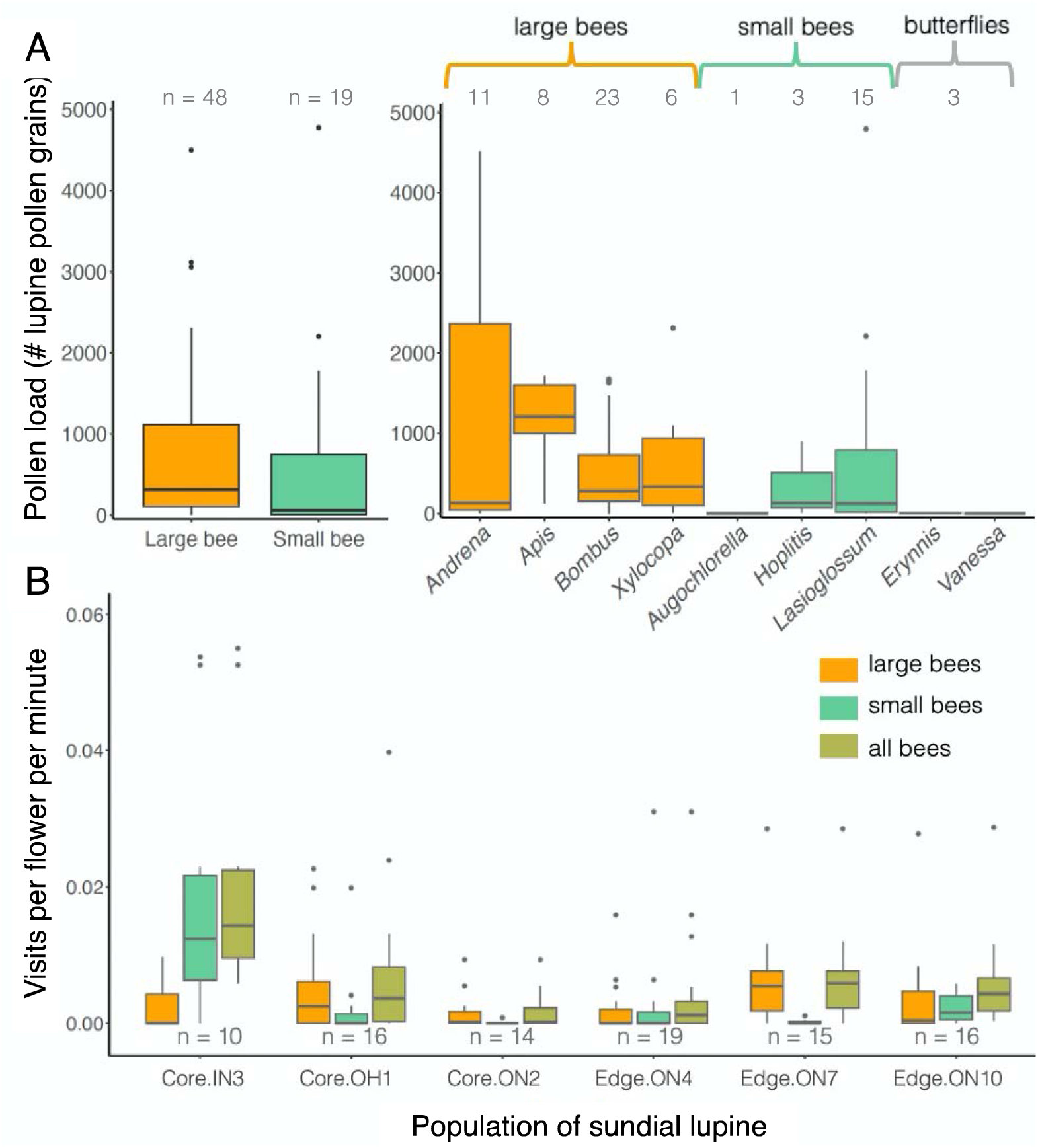
Visitation rates and visitor pollen loads. A) Pollen loads from all visitors seen visiting lupine flowers over 2 wk in population Core.ON2 in 2023; we saw four genera of large bees (orange), three genera of small bees (teal), and two butterfly genera (grey) visiting lupine flowers. Pollen loads carried by large and small bees did not differ significantly (Table S3). Numbers along the top indicate the number of bees sampled. **B)** Visitation rates of large bees, small bees, and all bees seen during 1084 min of observation in 2022. n = observation periods (10 to 20 min) per population. Bars show raw data: horizontal lines, boxes, and whiskers show the median, 1^st^ and 3^rd^ quartiles, and the 1.5× interquartile range above, respectively, while points show outliers.

**Table 1.** Pollen deposited during single visits to virgin lupine flowers. . Each visitor is an individual bee and each row is a different flower (6 bees visited a total of 15 flowers on 6 plants). Flowers were bagged except during the single visit, and all pollen deposited on each stigmas was counted. The second *Lasioglossum* bee evaded capture so can only be identified to genus.

| Visitor | Pollen grains deposited per stigma in 1 visit |  |  |
| --- | --- | --- | --- |
| | # Lupine | # Non-lupine | Mean ± SE Lupine |
| *Bombus impatiens* | 37 | 6 |  |
|  | 57 | 8 |  |
|  | 11 | 3 |  |
| | 25 | 11 | $32.5 \pm 9.7$ |
| *Bombus impatiens* | 11 | 0 |  |
| | 9 | 0 | $10.0 \pm 1.0$ |
| *Bombus bimaculatus* | 64 | 2 |  |
| | 7 | 1 | $35.5 \pm 28.5$ |
| *Xylocopa virginica* | 67 | 6 |  |
|  | 23 | 0 |  |
|  | 1 | 2 |  |
|  | 20 | 2 |  |
| | 15 | 1 | $25.2 \pm 11.1$ |
| *Lasioglossum vierecki* | 0 | 0 | $0 \pm \text{NA}$ |
| *Lasioglossum* sp. | 1 | 0 | $1 \pm \text{NA}$ |

### Visitation rates

During 1084 min of observation across 118 observation periods at six populations in 2022, we saw 267 bees and one butterfly visit lupine flowers. The only large bees seen in 2022 were from genera *Bombus* and *Xylocopa,* and they were the most common visitors observed (134 bees observed).

Visitation rates to lupine were fairly low (Fig. 3B). Overall visitation rates (all bees) differed among populations (*site*: χ^2^ = 29.9, *df* = 5, *P* < 0.0001), but not between the range core and edge (*range position*: χ^2^ = 0.2, *df* = 1, *P* = 0.69). The relative visitation rates for large and small bees differed among sites (*bee type x site*: χ^2^ = 41.2, *df* = 5, *P* < 0.0001), primarily because visitation rates by small bees were high in population Core.IN3. For large bees, the confirmed pollinators, visitation rates did not differ significantly between sites. For small bees, visitation rates were higher in Core.IN3 than any other site. Site Core.ON2 had lower visitation rates than Edge.ON4 and Edge.ON10, and Edge.ON10 had higher visitation rates than Edge.ON7.

### Pollen receipt

Lupine flowers received from 0 to 68 pollen grains on their stigma. Pollen receipt did not differ systematically between range-core and range-edge sites (*range position*: χ^2^ = 43.9, *df* = 5, *P* < 0.0001), but did vary significantly among sites (*site*: χ^2^ = 43.9, *df* = 5, *P* < 0.0001; Fig. 4A). Plants in population Core.OH1 received significantly more pollen than plants in most other populations, while plants in Edge.ON10 received less than most other populations. Pollen receipt did not vary significantly among collection dates during 2 weeks of peak flowering in Core.ON2 (*date*: χ^2^ = 3.5, *df* = 1, *P* = 0.062; Fig. S4).

**Figure 4.**
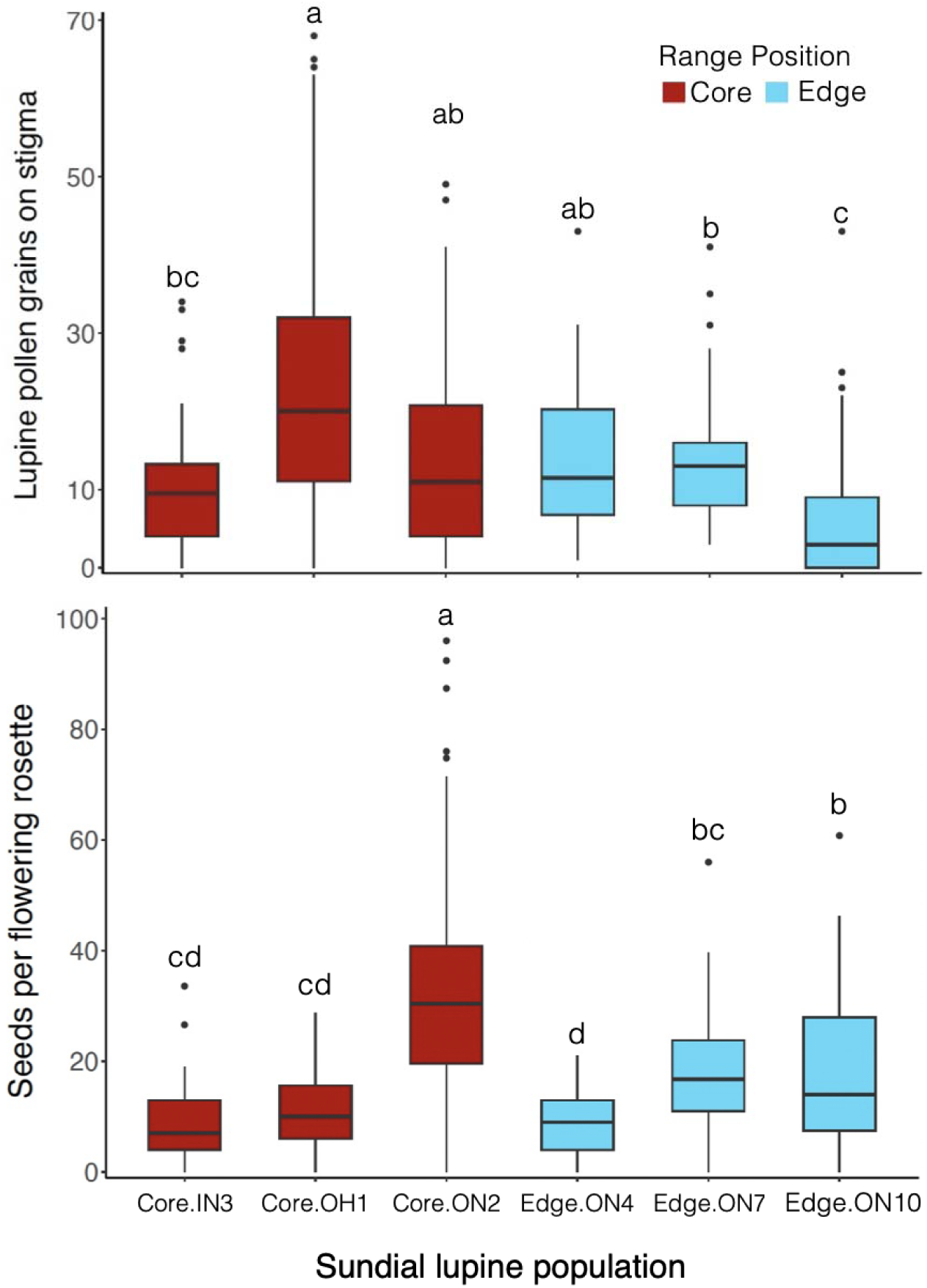
Pollen receipt and seed production in populations of sundial lupine at the species’ range-core or northern range edge. Neither pollen receipt (**A**; measured in 2023) nor seed production (**B**; measured in 2022) differed among range positions, but both differed among sites (statistical results in text). Boxplot formatting as in Fig. 3. Within each panel, boxes that do not share a letter are significantly different from each other.

### Seed production

Flowering rosettes produced 0 to 96 seeds (Fig. 4B). Seed production per flowering rosette did not vary between range positions (*range position*: χ^2^ = 0.05, *df* = 1, *P* = 0.82), but did vary among sites (*site*: χ^2^ = 78.7, *df* = 5, *P* < 0.0001). Flowering rosettes in Core.ON2 produced more seeds than any other site, followed by rosettes in Edge.ON7 and Edge.ON10. At the population level, mean seeds/flowering rosette was not correlated with mean visitation rate by all bees (χ^2^ = 2.9, *df* = 1, *P* = 0.089) or by large bees ( ^2^ = 0.2, *df* = 1, *P* = 0.66). Indeed, the population with the highest seed production (Core.ON2; Fig. 4B) had the lowest mean visitation rates (Fig. 3B).

## Discussion

We did not find evidence that declining sexual reproduction limits the northern range of sundial lupine. In 2022, seed production was highest at a core population and lowest at an edge population, but did not differ much among the remaining four populations, and was not lower in edge vs. core populations overall (Fig. 4B). Similarly, pollen receipt was highest at a core population and lowest at an edge population, but did not differ among the remaining four populations nor among range positions (Fig. 4A). Finally, while overall visitation rates to lupine flowers were highest in a core population, visitation rates by large bees, the only confirmed effective pollinators, were similar across populations (Fig. 3B). Our study therefore adds to a growing body of literature suggesting that pollination is not a key range-limiting factor across plant species. While biologists have recently recognized that the role of biotic interactions (Wisz et al. 2013) in limiting species ranges is under-studied, particularly for mutualisms (Paquette and Hargreaves 2021, Stephan et al. 2021, Fowler et al. 2023), there is no evidence to date that pollination is a particularly important constraint, more so than any other ecological factor which contributes to some range limits but not others (Dawson-Glass and Hargreaves 2022).

We also did not find evidence that pollination limits seed production in general for sundial lupine, though we were unable to test this directly as our pollen-supplementation experiment failed due to flower abortion. Neither overall visitation rates nor visitation rates by large bees correlated with seed production at the population level in 2022. Nor were populations with the highest visitation rates (Core.IN3) or seed production (Core ON2) in 2022 the populations with the highest pollen receipt (Core.OH1) in 2023. Equally, the populations with the lowest visitation (Core.ON2), were not those with the lowest seed production (Edge.ON4) or pollen receipt (Edge.ON10). While we might have found stronger relationships between visitation rates and seed production if we had measured them on the same plants, or between these metrics and pollen receipt if we had measured all metrics in the same year, the lack of correlation across plants and years suggests that at the population level, sexual reproduction is not chronically limited by lack of pollination. This suggests that lower reproductive success of some sundial lupine populations (Fig. 4) is caused by factors other than pollination.

One possible ecological reason why pollination may be decoupled from seed production in sundial lupine is that seed production may be limited by resources, rather than pollen receipt (Burd 1994). This could include lack of nutrients or light, inadequate climate (Pavlovic and Grundel 2009). This hypothesis warrants further testing, both with further pollen supplementation experiments, which are the classic tool for distinguishing pollen vs. resource limitation (Knight et al. 2005), and with independent assessments of habitat quality vs. seed production. To the extent that site quality is reflected in population size, we found mixed evidence that seed production varies with site quality; seed production was lowest in the smallest population (Edge.ON4) and highest in the largest population (Core.ON2), but did not vary consistently with size among the other four populations (Table. S1).

A second ecological explanation for the mismatch between pollination and seed production is that seed set may be limited by the quality of pollen received, rather than by the amount (Aizen and Harder 2007). Sundial lupine experiences limited self-compatibility and significant early inbreeding depression (Shi et al. 2005, Michaels et al. 2008), thus self-pollen is not nearly as beneficial for seed production as outcross pollen. While large bees transferred pollen successfully, they tended to visit all open flowers on a rosette. This might facilitate geitonogamy (self-pollination among flowers on the same plant), though this should be mitigated by their tendency to visit female flowers before male flowers. Testing for pollen-quality limitation would require pollen-supplementation experiments using both selfed and outcrossed pollen (Aizen and Harder 2007), or more single-visit experiments that track not only pollen deposition, but pollen tube growth or, ideally, seed production.

Our conclusions about which bees are effective pollinators differ somewhat from previous studies that were based on observation alone. We found that large bees, especially *Bombus* and *Xylocopa*, were the sundial lupine’s most effective pollinators, as previously deduced based on observations of bee foraging (Bernhardt et al. 2008, Petitta et al. 2024). However, previous observational work also categorized small bees, particularly in the genus *Osmia* (mason or orchard bees), as effective pollinators (Bernhardt et al. 2008, Petitta et al. 2024). Neither our observations nor single-visit experiment supported this conclusion; we rarely saw small bees manipulate flowers in a way that would lead to stigma contact, and small bees transferred almost no pollen in single visit experiments. As there is often a mismatch between animals that visit flowers and those that pollinate (King et al. 2013), particularly for plants that offer pollen as a reward (Thomson and Goodell 2001, Hargreaves et al. 2009), experimental confirmation of pollinator effectiveness is important. Nevertheless, we did not observe *Osmia* bees at our six sites, and our sample size for single-visit experiments was low for small bees. Thus further such experiments would be needed to confirm whether small bees, particularly *Osmia*, effectively pollinate sundial lupine.

Finally, the relatively low visitation rates we observed may be typical for sundial lupine, despite its striking floral displays. Across our 6 sites we observed 0.25 visitors/min/m^2^; the only other quantification of visitation rates to sundial lupine saw even fewer visitors across 10 sites (271 visitors during 4170 plot-observation minutes on 1×2 m plots, i.e. 0.03 visitors/min/m^2^) (Bernhardt et al. 2008). Few other studies have quantified visitation rates to lupine species, but visitation rates to sundial lupine are comparable to those observed for *Lupinus arboreus,* another pollen-rewarding species, in both its native and exotic range (∼0.031 and 0.014 bee visitors/min/raceme)(Stout et al. 2002, Rivest et al. 2024). Despite the low visitation rates, sundial lupine may still be a critical source of pollen (i.e. protein), for the native bee populations in its range.

## Conclusion

We found no evidence that pollination chronically limits seed production of sundial lupine, nor evidence that low pollination or seed production limit the species’ northern range. The most fruitful avenues of future research for lupine’s conservation would be: a) further pollen addition and single visit experiments to test whether seed production is limited by pollen quality, b) further single-visit experiments to test whether small bees, particularly *Osmia*, transfer pollen, and c) surveys or transplant experiments to test which habitat components and life-stages limit population success, particularly at the species’ northern range edge.

## Author Contributions

This work comes from MB’s MSc thesis, supervised by ALH. ALH and MB designed the study. MB collected the data with help from CPS, performed analyses and wrote the first draft of the manuscript with guidance from ALH. Both authors contributed to analyses and writing of the final version, all authors edited the final version.

## Data Availability

All data and code will be deposited in the Borealis Digital Repository upon submission.

## Acknowledgements

We declare no conflict of interest. We thank Nicole Cappellazzo, April Kowalchuk-Reid and Andrew Kemp for help in the field, and Dr. Helen Michaels and Mary Gartshore for sharing their extensive knowledge about lupine. We also thank Ontario Parks, the Nature Conservancy of Canada, the City of Toronto, the Indiana Department of Natural Resources, the Nature Conservancy at Indiana, Metroparks Toledo, and Rick Scott for ranting permission and facilitating access for fieldwork. We thank all the bees and plants that contributed to this work. This work is part of a NOVA Alliance grant to ALH ‘Using range-edge ecology to inform conservation of Canada’s rare plants’ from the Natural Sciences and Engineering Research Council of Canada (NSERC) and Fonds de Recherche du Québec - Nature et Technologies (FRQNT). It was also funded through an NSERC Discovery Grant to ALH, FRQNT scholarship to MB, an NSERC Canada Graduate Scholarship to CPS, Canadian Foundation for Innovation JELF grant to ALH, NSERC undergraduate research award to AK and McGill undergraduate research award to NC.

## Supplementary Material

### Supplementary Figures and Tables

**Figure S1.**
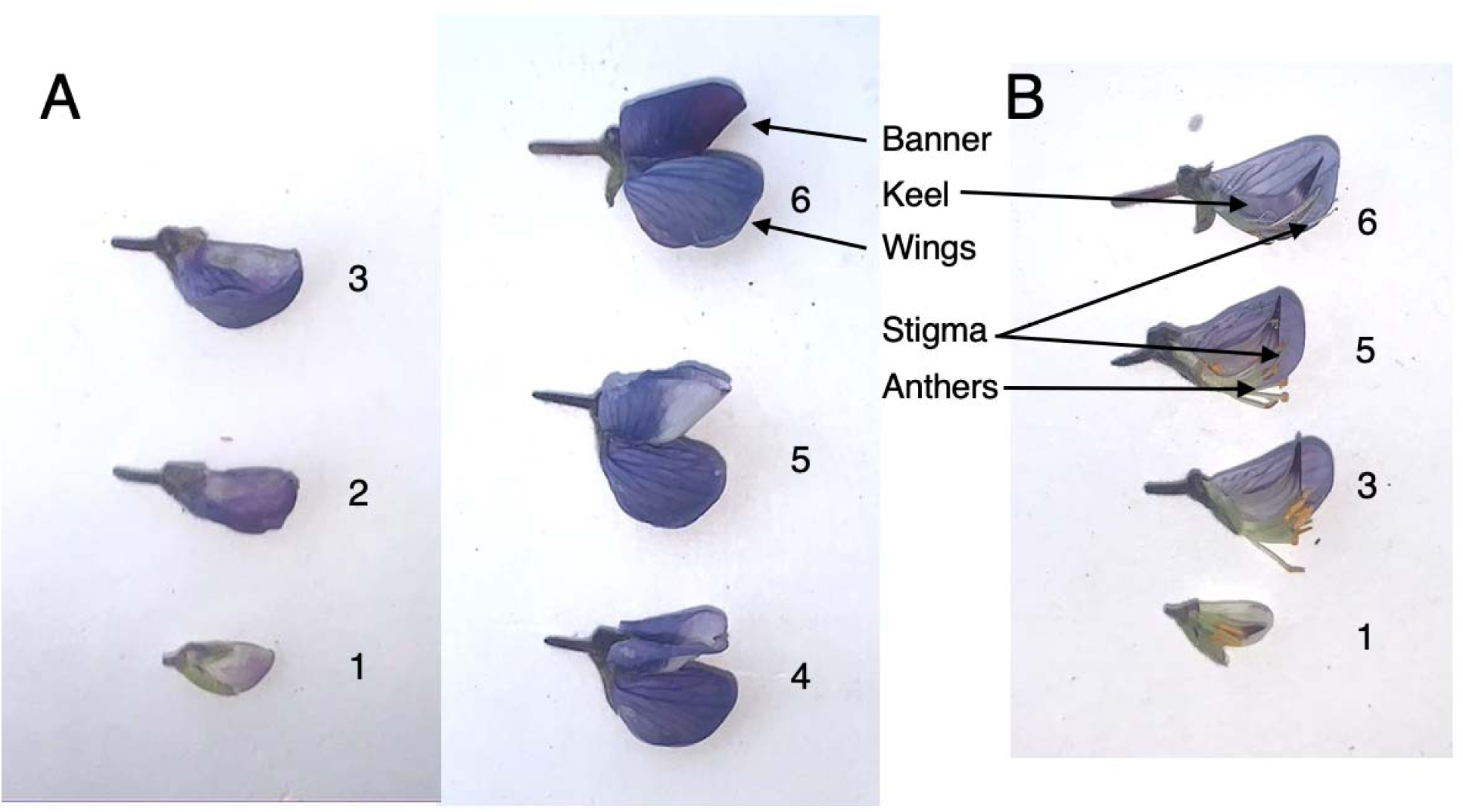
Development of sundial lupine flowers. Panels show external (**A**) and internal (**B**) morphology. In both panels, numbers indicate the flower’s stage: 1) young bud, 2) older bud, 3) flower is just about to open and begin producing pollen, 4) flower is producing pollen but stigma is not elongated or receptive (receptivity judged by stigma appearance), 5) stigma is elongated and receptive, 6) flower is wilting and no longer receptive. Flowers are open to pollinators (stage 4 to 5) for 3 to 4 days.

**Figure S2.**
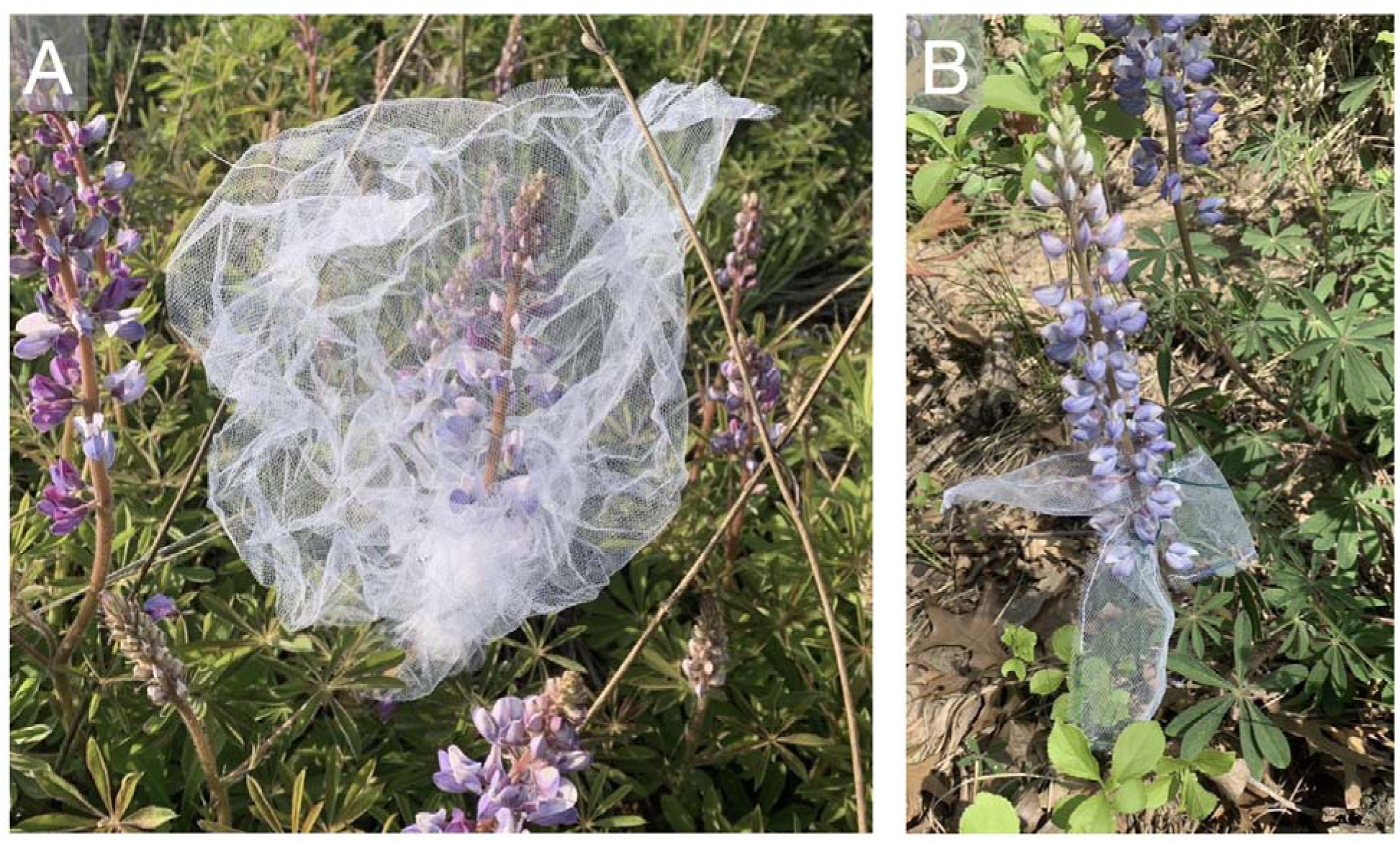
Tulle bags used to exclude pollinators. **A**) 40 x 30 cm bag around an entire inflorescence, secured around the base of an inflorescence with a twist tie. **B**) Three 15 x 8 cm bags on individual flowers, secured around the base of each flowering stalk with a twist tie. While large bags like that in A did not seem to affect flowers or inflorescences for the duration we deployed them (2 to 4 days), all individually bagged flowers like those in B aborted before producing seed.

**Figure S3.**
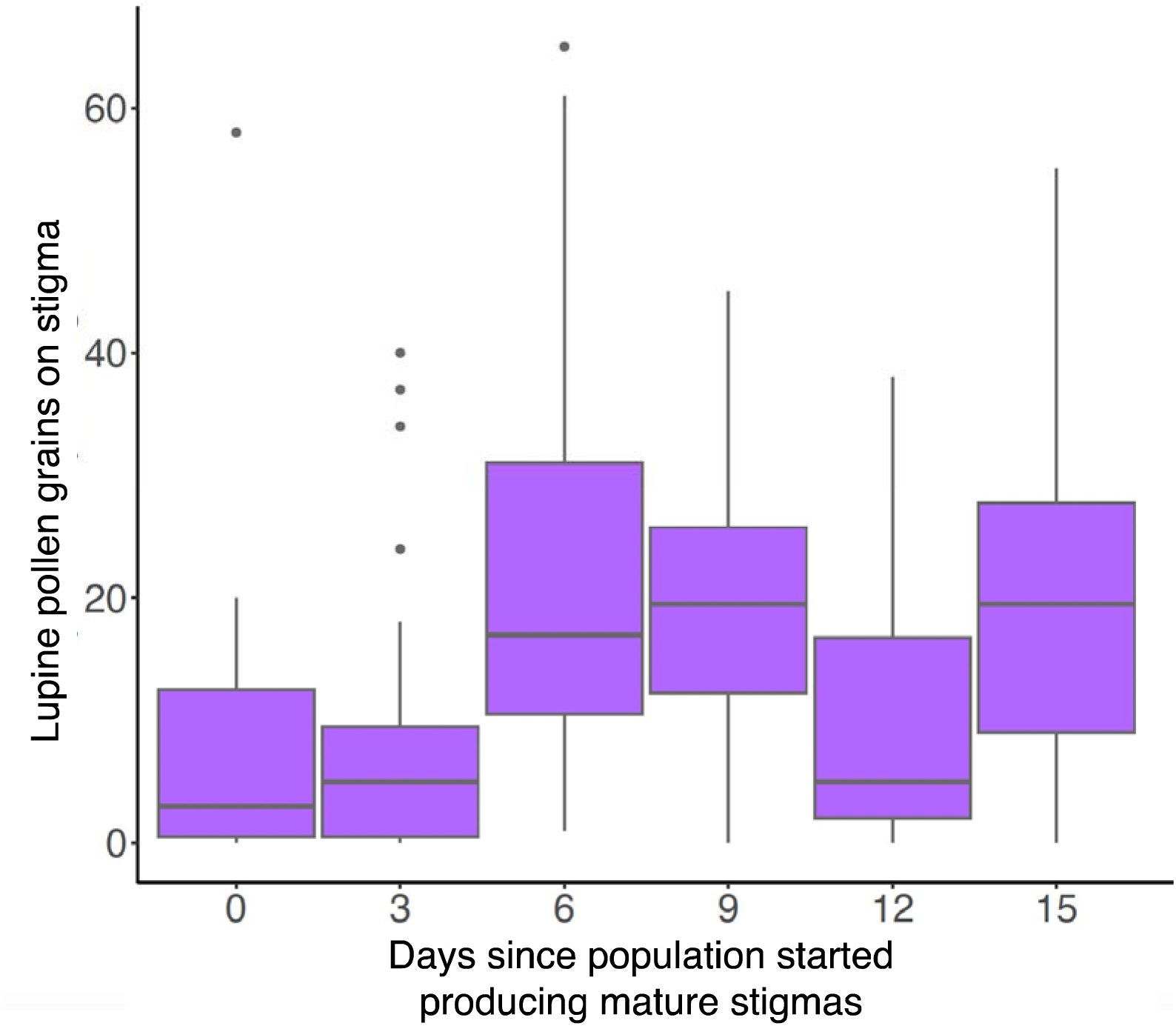
Pollen receipt through time in population Core.ON2. Data were collected over 2 weeks during peak flowering in 2023. Stigmas were collected every 3 days starting on Day 0, which was the first day that receptive stigmas were seen in the population. Pollen receipt did not differ significantly among collection dates. Bars show raw data: horizontal lines, boxes, and whiskers show the median, 1^st^ and 3^rd^ quartiles, and the 1.5× interquartile range above, respectively, while points show outliers.

**Table S1.** Population details. Populations names (ordered by latitude) denote: whether it was considered to be in sundial lupine’s range core or at its northern range edge; the state or province the population is in, and a unique identifying number (numbers are not sequential because they are from a larger study with additional populations; *forthcoming*). Because it is hard to reliably distinguish genetically distinct individuals of sundial lupine, population sizes are the number of lupine clusters rather than individuals *per se*. They indicate the relative sizes of the populations, rather than their absolute sizes. Each cluster is a group of lupine rosettes, up to 2’ diameter, that share similar phenology and leaf morphology.

| Population | Location | Latitude | Longitude | Size class<br>(1000s of lupine<br>clusters) |
| --- | --- | --- | --- | --- |
| Core.OH1 | Lou Campbell Tall Grass Prairie | 41.535 | -83.837 | 10 – 15 |
| Core.IN3 | Indiana Dunes State Park | 41.662 | -87.051 | 1 – 5 |
| Core.ON2 | Norfolk County Backus Woods | 42.682 | -80.476 | 20 – 30 |
| Edge.ON4 | Pinery Provincial Park (forest) | 43.234 | -81.861 | .1 – .5 |
| Edge.ON7 | High Park Central | 43.647 | -79.467 | .5 – 1 |
| Edge.ON10 | Rice Lake Plains Lilac Valley | 44.136 | -78.142 | 1 – 5 |

**Table S2.** Pollen loads. Data include all insects seen visiting sundial lupine flowers at site Core.ON2 during 2 wk of peak flowering in 2023. n is the individuals swabbed for pollen.

| Visitor functional group | Lupine pollen grains carried per visitor type |  |  |
| --- | --- | --- | --- |
| Taxonomy | Mean | Min – Max | n visitors |
| Large bees (n = 48 individuals) |  |  |  |
| Apidae |  |  |  |
| Apis |  |  |  |
| *Apis mellifera* | 1167 | 128 – 1707 | 8 |
| Bombus |  |  |  |
| *Bombus bimaculatus* | 768 | 25 – 1667 | 11 |
| *Bombus impatiens* | 290 | 1 – 845 | 9 |
| *Bombus pensylvanicus* | 227 | 71 – 282 | 3 |
| Xylocopa |  |  |  |
| *Xylocopa virginica* | 691 | 13 – 2304 | 6 |
| Andrenidae |  |  |  |
| Andrena |  |  |  |
| *Andrena carlini* | 1024 | 0 – 4500 | 8 |
| *Andrena crataegi* | 10 | – | 1 |
| *Andrena miserabilis* | 5 | – | 1 |
| *Andrena spp.* | 3117 | – | 1 |
| Small bees (n = 19 individuals) |  |  |  |
| Halictidae |  |  |  |
| Augochlorella |  |  |  |
| *Augochlorella aurata* | 1 | – | 1 |
| Lasioglossum |  |  |  |
| *Lasioglossum imitatum* | 198 |  | 1 |
| *Lasioglossum vierecki* | 346 | 3 – 827 | 6 |
| *Lasioglossum spp. I* | 4778 |  | 1 |
| *Lasioglossum spp. II* | 892 | 5 – 1779 | 2 |
| *Lasioglossum spp. III* | 753 | 5 – 2203 | 3 |
| *Lasioglossum spp. IV* | 4 | 0 – 8 | 2 |
| Megachilidae |  |  |  |
| *Hoplitis pilosifrons* | 346 | 11 – 892 | 3 |
| Butterflies (n = 3 individuals) |  |  |  |
| Erynnis |  |  |  |
| *Erynnis spp.* | 7 | 1 – 12 | 2 |
| Vanessa |  |  |  |
| *Vanessa atalanta* | 3 | – | 1 |

### Supplementary Methods: Breeding System and Pollen Limitation Experiments

We attempted to quantify self compatibility, capacity for autonomous self-pollination, and pollen limitation in each of our six populations, using the methods below. However, all individually-bagged flowers aborted, so no data are available from these experiments.

#### Breeding system

On the first day we visited each population, we selected 30 inflorescences, at least 2 m apart from each other, that had flowers about to open and no mature (i.e. already receptive) flowers. We isolated each from pollinators by enclosing the entire inflorescence in a 45 x 30 cm white tulle bag secured at the base of the inflorescence using a twist tie (Fig. S2A). Bags were left on the inflorescence for two days, until some flowers were receptive. On the third day, we removed the bags and selected three flowers that appeared receptive and assigned them randomly to one of three treatments: hand pollinated with pollen the same inflorescence (selfed), hand-pollinated with pollen from an inflorescence at least 5 m away (outcrossed), or unmanipulated to test for autonomous self-fertilization (autogamous). Pollen was applied to the stigma using tweezers, as pollen stuck together in large chunks; we were carefully not to touch the metal tweezers to the stigma surface to avoid damage, and tweezers were dipped in ethanol and dried between plants and pollen donors. Pollen for outcrossing was collected on the morning of the experiment. We then bagged each flower individually with a 15 x 8 cm white tulle bag secured with twist ties around the pedicle, to capture seeds if pods burst before we returned. Flowers and bags were left until seed set.

#### Pollen limitation

To test whether seed production was limited by the quantity of pollen received, we conducted pollen supplementation experiments. We selected an additional 30 inflorescences, (at least 2 m apart from each other and from the breeding-system plants when possible), that were young and had some early female-phase flowers. We selected two flowers per inflorescence, assigned one as a control (unmanipulated) and supplemented the other with outcross pollen at least once, but often once on the first day and again the following day. We then bagged each flower individually as in the breeding system experiment.

